# Pharmacokinetics and *In vivo* Efficacy of Epetraborole against *Burkholderia pseudomallei*

**DOI:** 10.1101/2025.07.10.664055

**Authors:** Jason E. Cummings, Danara Flores, Vincent E. Guglielmi, Gregory Dooley, M. R. K. Alley, Richard A. Slayden

## Abstract

Melioidosis, caused by *Burkholderia pseudomallei*, remains a major therapeutic challenge due to high relapse rates and intrinsic antibiotic resistance. Epetraborole (EBO), a leucyl-tRNA synthetase inhibitor, represents a novel therapeutic approach with a distinct mechanism of action compared to standard-of-care antibiotics. Preclinical studies included MIC determination, pharmacokinetic (PK) profiling, dose range and fractionation studies, and efficacy assessments in a 24-hour post-bacterial challenge model of a murine *B. pseudomallei* lung infection. EBO demonstrated a clear dose-dependent reduction in lung bacterial burden. Doses ≥200 mg/kg (achieving AUC₀–₂₄ ∼110 µg·h/mL) produced >1.6 log₁₀ CFU decreases from the start-of-therapy baseline across all ten *B. pseudomallei* strains. Notably, an AUC₀–₂₄ of ∼110 µg·h/mL was achieved in humans with a 2000 mg IV dose in a phase 1 clincial trial where doses up to 4000 mg per day for 14 days were well tolerated with no serious adverse events or dose-limiting adverse events. When EBO doses of 600, 300 and 100 mg/kg SC were fractionated by once, twice and three times a day against the *B. pseudomallei* strain NCTC7383, which represents the MIC_100_ strain, the efficacy indicated that the pharmacokinetics-pharmacodynamics (PK/PD) driver of epetraborole is total drug exposure (AUC) rather than peak concentration (Cmax) or time above MIC. The inhibition of leucyl-tRNA synthetase represents a unique molecular target, reducing cross-resistance potential with existing β-lactam antibiotics and enabling combination therapy strategies. These findings substantiate EBO as a promising therapeutic option for clinical melioidosis to improve treatment outcomes. Notably, this study represents the first demonstration of *in vivo* efficacy against a panel of ten genetically and geographically diverse *B. pseudomallei* strains in a murine model. This unprecedented breadth of strain coverage provides strong evidence of EBO’s robust and strain-independent therapeutic potential.

**Author Summary:** Melioidosis is a life-threatening disease caused by *Burkholderia pseudomallei*, a bacterium found in soil and water in tropical regions. It is challenging to treat because it is often resistant to standard antibiotics, requires long courses of therapy, and can relapse even after treatment. New treatment options are urgently needed. Epetraborole is an experimental antibiotic with a novel mechanism that blocks bacterial protein synthesis. In this study, we tested how well epetraborole works in mice infected with *B. pseudomallei*, using ten different strains of the bacteria collected from geographically diverse regions. This is the first study to test a melioidosis treatment against such a broad panel of *B. pseudomallei* strains in an animal model. We found that epetraborole significantly reduced bacterial counts in the lungs of infected mice. The antibiotic remained effective even against strains with higher resistance and high bacterial loads 24 hours after infection, showing promise for severe clinical infections. Our findings suggest that epetraborole could be a valuable addition to existing melioidosis treatments, especially in regions with limited therapeutic options.

## Introduction

*Burkholderia pseudomallei*, a Gram-negative bacterium, is the causative agent of melioidosis, a severe tropical infectious disease with a significant threat to public health and a nationally notifiable condition in the US[1]. Melioidosis is endemic in tropical regions, particularly Southeast Asia and Northern Australia, where *B. pseudomallei* is found in soil and water. Clinical manifestations of melioidosis range from localized skin or soft tissue infections to severe pneumonia and septicemia, often leading to high mortality rates. Due to its diverse clinical presentations, resemblance to other tropical diseases, and the potential for chronic relapses, melioidosis poses diagnostic challenges, and its epidemiology remains underreported in many endemic regions. Cases have emerged in non-endemic regions due to epidemiological transitions, zoonotic hazards, and climate change[2, 3]. The clinical significance of *B. pseudomallei* is heightened by its intrinsic resistance to various antibiotics, making treatment challenging and necessitating prolonged and combination therapies. Novel treatment options that can be used alone or in combination are critical for addressing the clinical burden of melioidosis, emphasizing the importance of drug discovery research efforts to combat this neglected tropical disease[4, 5].

The clinical management and treatment of *B. pseudomallei* involve a combination of antimicrobial therapy and supportive care. IV Ceftazidime or meropenem monotherapy are the first-line antibiotics for the current standard of care (SoC) treatment of severe melioidosis, while oral trimethoprim-sulfamethoxazole is employed for the longer-term eradication therapy (12-20 weeks). Combination therapy enhances efficacy and reduces the risk of resistance development, and the duration of treatment varies based on the severity of the infection and the patient’s response. In some cases, prolonged treatment courses and a step-down approach with oral antibiotics may be necessary for complete eradication. Long-term antimicrobial therapy is required to prevent relapse, complicating disease management[6]. Supportive care measures, including respiratory and hemodynamic support, are essential for managing side effects associated with prolonged treatments for severe cases. The management of melioidosis requires early initiation of appropriate antimicrobial therapy, which is crucial for improving patient outcomes[5, 7, 8].

*B. pseudomallei* poses a significant challenge in antimicrobial therapy due to its intrinsic resistance to various antibiotics. This bacterium exhibits resistance or tolerance to a broad spectrum of antimicrobial agents, including beta-lactams, aminoglycosides, and macrolides, limiting treatment options[9]. Moreover, transient resistance and the ability of *B. pseudomallei* to form biofilms during infection further enhance its resistance to antibiotics, hindering the penetration of drugs and promoting chronic infections[10]. The emergence of multidrug-resistant strains has raised concerns about treating and managing melioidosis [11]. Studies have highlighted the complex mechanisms underlying drug resistance in *B. pseudomallei*, including efflux pumps, cell permeability changes, and genetic mutations. The limited availability of effective antibiotics underscores the urgent need to develop novel therapeutic strategies to combat drug-resistant strains of *B. pseudomallei* [12–14]

Epetraborole (EBO), a leucyl-tRNA synthetase (LeuRS) inhibitor, is a broad-spectrum investigational drug that disrupts protein synthesis, leading to bacterial cell death. In preclinical studies, EBO has demonstrated potent activity against Gram-negative bacteria, including *B. pseudomallei*. EBO’s efficacy against *B. pseudomallei* suggests its potential as a therapeutic option for melioidosis, addressing the need for alternative treatments [15]. Further clinical trials are necessary to evaluate its safety and effectiveness in human subjects.

While most prior efficacy studies have relied on one or two well-characterized strains, such as 1026b or K96243, this study expands the scope by evaluating *in vivo* efficacy across ten distinct *B. pseudomallei* isolates[16]. This is the first known preclinical therapeutic study to assess drug performance across such a broad strain panel, thereby enhancing the translational value of the findings [9]. The breadth of this study aligns with a US FDA recommendation that for *B. pseudomallei*, 10 strains are studied for robust PK/PD target attainment and between-strain variability characterization (SD Prior Personal Communication to the authors). This assessment of EBO includes pharmacokinetic analysis, dose fractionation, dose response, and efficacy in an acute animal model of melioidosis designed to look at bactericidal activity over 48 hours of therapy after 24 hours of infection. These data demonstrate that advanced exploration of EBO in the therapy of melioidosis is warranted, further supporting LeuRS as a clinically relevant drug target.

## Materials and Methods

### Ethics statement

All studies performed at Colorado State University were conducted in a BSL3 facility dedicated to bacterial pathogen work under the approvals and management of the Biosafety Official. Studies were approved by the Institutional Biosafety Committee and the Institutional Animal Care and Use Committee and performed under approvals PARF 17-095B and IACUC protocol 3796.

### Inoculum size effect on minimum inhibitory concentration (MIC) determination

Reference strains (*B. pseudomallei* 1026b, MSHR435, NCTC7383) were used based on prior susceptibility data [15]. For each evaluation, bacteria were prepared fresh by growth from the standard Luria-Bertani (LB) agar stocks at 37°C for 48-72 hours. Bacteria recovered from the LB plates were used to inoculate 10 mL LB broth. Broth cultures were then incubated for 18 hours at 37°C, diluted 1:100, and incubated for another 6 hours at 37°C. Bacteria were then diluted to final concentrations of ∼1×10^8^, 1×10^7^, 1×10^6^, 1×10^5^, and 1×10^4^ CFU/mL concentration in cation-adjusted Mueller-Hinton (caMH) Broth (MilleporeSigma, St. Louis, MO) and added to each well for each drug plate for a final 1:2 dilution. The concentration range tested for epetraborole, ceftazidime, meropenem, doxycycline, and chloramphenicol was 0.03 – 64 µg/mL in caMH broth. MIC plates were incubated at 37°C for 18 hours, at which time MIC was determined per CLSI guidelines (CLSI, 2018). All drug stocks were validated by MIC determination against strains *E. coli* ATCC 25922 and *P. aeruginosa* ATCC 27853, and values were compared to published values.

### Drug Exposure Pharmacokinetic Analysis

Female BALB/c mice (7–9 weeks old) were dosed subcutaneously with epetraborole at 30, 100, 200, 300, 400, or 600 mg/kg. Blood was collected by terminal cardiac puncture at 0.5, 1.5, 3, 6, 12, and 24 hours (n = 3/timepoint). Blood was placed into K2EDTA tubes, mixed well, and centrifuged at 2,000xg, 4 ^°^C for 10 minutes. 40μl plasma was mixed with 120μl 75% v/v methanol: water (LC-MS grade), vortexed to mix for 10 seconds, and stored at -20°C until ready for mass spectrometry analysis. Samples were analyzed via LC/MS/MS to determine the concentration of epetraborole within mouse plasma. Sample concentrations were determined by creating a 2-fold standard curve with 10 calibration standards (2 to 1024 ng/mL) prepared fresh in mouse plasma and a blank. Samples and calibrants underwent protein precipitation via cold methanol extraction, followed by a spike of 20ng of internal standard. Once analyzed, samples out of range were diluted in Burdick & Jackson HPLC Water. The chromatographic samples and calibrants were injected into a reversed-phase HPLC column (Waters Atlantis T3 column maintained at 25°C). Epetraborole and internal standard were detected in positive electrospray ionization in the multiple reaction monitoring mode on an Agilent 6460 triple quad mass spectrometer. A gradient from 100% water to 100% methanol was used to run and clean the column after each sample.

### Lab reference & clinical strain infection comparison of thirteen *B*. pseudomallei strains in an animal infection model

Acute infection disease profiling of *B. pseudomallei* strains 1026b, MSHR435, NCTC7383, K96243, 406e, NCTC6700, HBPUB10134a, HBPUB10303a, NCTC10274, NCTC10276, 1710a, 1710b, and 1106b was conducted in mice infected intranasally with 5,000 CFU. N=5 mice per group were euthanized at 24 hours to determine lung burden. The 24-hour timepoint was chosen as the optimal delayed dosing starting point for future studies due to the short duration of these acute studies (72 hours to morbidity). Mice were necropsied to harvest lungs and tissues, homogenized, diluted serially 1:10 in saline, and cultured onto LB agar to determine bacterial burden.

### Epetraborole dose fractionation study

To evaluate the *in vivo* efficacy of varying doses and dose frequency of epetraborole, mice were infected intranasally with 5,000 CFU *B. pseudomallei* NCTC7383, representing the EBO MIC_100_ for *B. pseudomallei*. A total of 4 mice were sacrificed 24 hours post-infection to determine the pre-treatment bacterial burden in the lung. Starting at 24 hours post-infection, mice were treated with EBO subcutaneously at 600 mg/kg once daily (QD), 300 mg/kg BID twice daily (BID), 200 mg/kg three times per day (TID), 300 mg/kg QD, 150 mg/kg BID, 100 mg/kg TID, 100 mg/kg QD, 50 mg/kg BID, and 33 mg/kg TID. A standard-of-care control group received ceftazidime TID subcutaneously at 300mg/kg. All mice received treatment for 48 hours (72 hours post-infection). At 72 hours post-infection, the remaining mice were necropsied to harvest lungs, tissues were homogenized, diluted serially 1:10 in saline, and cultured onto LB agar to determine bacterial burden.

### Epetraborole dose response against ten *B. pseudomallei* strains

To better assess the *in vivo* efficacy of epetraborole, a dose-response efficacy study was performed against lab reference and clinical strains 1026b, MSHR435, NCTC7383, K96243, 406e, HBPUB10134a, HBPUB10303a, 1710a, 1710b, and 1106b. Mice were infected intranasally with 5,000 CFU, and N=5 mice per group were euthanized at 24 hours to determine lung burden. At 24 hours post-infection, mice were treated with epetraborole subcutaneously at 200 mg/kg, 100 mg/kg, 30 mg/kg, 10 mg/kg, and 3 mg/kg once daily for 48 hours (72 hours post-infection). At 72 hours post-infection, the remaining mice were necropsied to harvest lungs, and tissues were homogenized, diluted serially 1:10 in saline, and cultured onto LB agar to determine bacterial burden.

### Statistical analysis, effect size calculation, and performance classification

All statistical analyses were performed using GraphPad Prism or equivalent software. Differences in lung bacterial burden between treatment groups were assessed using two-way analysis of variance (ANOVA) followed by Tukey’s post-hoc multiple comparisons test. A p-value of <0.05 was considered statistically significant, with thresholds indicated as p < 0.05 (*), p < 0.001 (**), and p < 0.0001 (***) in figure panels. To quantify the magnitude of treatment effects, Cohen’s *d* effect sizes were calculated using the formula:

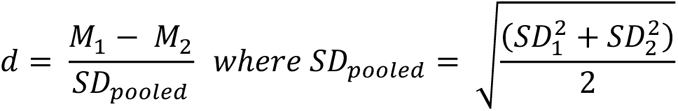

Here, M₁ and M₂ are the group means, and SD₁ and SD₂ are their respective standard deviations. For lung burden comparisons, M₁ and SD₁ correspond to the control group, and M₂ and SD₂ to the treatment group. We used a quartile-based approach specific to each dataset to classify effect sizes. The distribution of all calculated d values was divided into quartiles within each experiment (fractionation and efficacy studies separately). Effect size categories were then defined as minimal (1^st^ quartile), moderate (2^nd^ quartile), large (3^rd^ quartile), and very large (4^th^ quartile). This data-driven method allowed interpretation of effect sizes relative to the observed variability and biological response in each context. Sample sizes (n) for each group are provided in the figure legends and were selected based on prior experience with murine infection models.

## Results

### Effect of Inoculum Size on Minimum Inhibitory Concentrations (MICs)

The impact of bacterial inoculum size on antimicrobial susceptibility is a critical factor in determining the clinical efficacy of antibiotics and the standardization of MIC determinations. Therefore, we evaluated MIC values for EBO and comparator standard of care (SoC) drugs (ceftazidime, meropenem, doxycycline, and chloramphenicol) against *B. pseudomallei* strains 1026b, MSHR435, and NCTC7383 across a range of inoculum concentrations (10⁴ to 10⁷ CFU/mL; **Table 1**). Quality control strains *Escherichia coli* ATCC 25922 and *Pseudomonas aeruginosa* ATCC 27853 were included to validate assay performance.

**Table 1:** The Effect of Different Inoculum Concentrations on MIC (µg/mL) Values.

| Bacteria | Inoculum (CFU/mL) | EBO | Ceftazidime | Meropenem | Doxycycline | Chloramphenicol |
| --- | --- | --- | --- | --- | --- | --- |
| *B. pseudomallei* 1026b | $5 \times 10^7$ | >32 | >32 | >32 | >32 | >32 |
| *B. pseudomallei* 1026b | $5 \times 10^6$ | 1 | 4 | 2 | 0.5 | 8 |
| *B. pseudomallei* 1026b | $5 \times 10^5$ | 1 | 2 | 2 | 0.25 | 4 |
| *B. pseudomallei* 1026b | $5 \times 10^4$ | 0.5 | 2 | 1 | 0.25 | 4 |
| *B. pseudomallei* 1026b | $5 \times 10^3$ | 0.5 | 1 | 1 | 0.13 | 2 |
| *B. pseudomallei* MSHR 435 | $5 \times 10^7$ | >32 | >32 | >32 | >32 | >32 |
| *B. pseudomallei* MSHR 435 | $5 \times 10^6$ | 2 | 4 | 2 | 16 | 32 |
| *B. pseudomallei* MSHR 435 | $5 \times 10^5$ | 1 | 4 | 2 | 4 | 16 |
| *B. pseudomallei* MSHR 435 | $5 \times 10^4$ | 1 | 4 | 2 | 2 | 8 |
| *B. pseudomallei* MSHR435 | $5 \times 10^3$ | 1 | 2 | 2 | 1 | 8 |
| *B. pseudomallei* NCTC 7383 | $5 \times 10^7$ | >32 | >32 | >32 | >32 | >32 |
| *B. pseudomallei* NCTC 7383 | $5 \times 10^6$ | 4 | 2 | 1 | 4 | >32 |
| *B. pseudomallei* NCTC 7383 | $5 \times 10^5$ | 4 | 2 | 1 | 2 | >32 |
| *B. pseudomallei* NCTC 7383 | $5 \times 10^4$ | 4 | 2 | 1 | 2 | >32 |
| *B. pseudomallei* NCTC 7383 | $5 \times 10^3$ | 4 | 2 | 1 | 2 | >32 |
| *E. coli* ATCC 25922 | $5 \times 10^7$ | 2 | 8 | 8 | 1 | 8 |
| *E. coli* ATCC 25922 | $5 \times 10^6$ | 0.5 | 0.5 | 0.063 | 0.5 | 4 |
| *E. coli* ATCC 25922 | $5 \times 10^5$ | 0.5 | 0.13 | 0.031 | 0.5 | 4 |
| *E. coli* ATCC 25922 | $5 \times 10^4$ | 0.5 | 0.13 | 0.031 | 0.5 | 4 |
| *E. coli* ATCC 25922 | $5 \times 10^3$ | 0.25 | 0.13 | 0.031 | 0.5 | 4 |
| *P. aeruginosa* ATCC 27853 | $5 \times 10^7$ | >32 | >32 | >32 | >32 | >32 |
| *P. aeruginosa* ATCC 27853 | $5 \times 10^6$ | 2 | 2 | 0.5 | 16 | >32 |
| *P. aeruginosa* ATCC 27853 | $5 \times 10^5$ | 0.5 | 1 | 0.25 | 8 | >32 |
| *P. aeruginosa* ATCC 27853 | $5 \times 10^4$ | 0.5 | 1 | 0.25 | 8 | >32 |
| *P. aeruginosa* ATCC 27853 | $5 \times 10^3$ | 0.5 | 1 | 0.25 | 4 | >32 |

All *B. pseudomallei* strains exhibited a consistent increase in MIC values with rising inoculum. For strain 1026b, MICs at 5 × 10⁵ CFU/mL were 1 µg/mL for EBO, 2 µg/mL for ceftazidime, 2 µg/mL for meropenem, 0.25 µg/mL for doxycycline, and 4 µg/mL for chloramphenicol. At the highest inoculum (5 × 10⁷ CFU/mL), all five antibiotics had MICs exceeding 32 µg/mL. Strain MSHR435 followed a similar trend, with EBO MIC rising from 1 µg/mL to >32 µg/mL, and other antibiotics exhibiting ≥8- to >32-fold increases. NCTC7383 also showed inoculum-dependent MIC values, with EBO increasing from 1–4 µg/mL at standard inoculum to >32 µg/mL at high inoculum. This strain is resistant to chloramphenicol, so it was uniformly resistant to chloramphenicol (>32 µg/mL) at all inoculum levels.

Control strains showed expected MICs at 5 × 10⁵ CFU/mL: *E. coli* values ranged from 0.031–0.5 µg/mL for all tested drugs except chloramphenicol (4 µg/mL), while *P. aeruginosa* values ranged from 0.5–8 µg/mL. At 5 × 10⁷ CFU/mL, MICs increased to 1–8 µg/mL in *E. coli* and >32 µg/mL in *P. aeruginosa*, especially for β-lactams and doxycycline, confirming expected trends in inoculum-related resistance. Notably, while EBO MICs increased at high inocula, the magnitude of change was often less than that observed for ceftazidime or meropenem, especially at intermediate concentrations. This suggests a more stable potency profile for EBO under higher bacterial burden conditions.

### Pharmacokinetic analysis of Epetraborole

To determine the pharmacodynamic drivers for efficacy, we first evaluated the pharmacokinetics of EBO in satellite groups of BALB/c mice following subcutaneous (SC) administration at doses ranging from 30 mg/kg to 600 mg/kg. Plasma concentrations were measured at 0.5, 1.5, 3, 6, 12, and 24 hours post-dosing (**Figure 1**). A dose-dependent increase in plasma drug levels was observed, with peak concentrations (Cmax) occurring at 0.5 hours for all doses. At 30 mg/kg, the mean concentration at 0.5 h was 5.10 ± 0.47 µg/mL, whereas 600 mg/kg yielded a peak concentration of 68.23 ± 18.95 µg/mL. Drug levels declined over time, with substantial decreases by 6 hours and by 12 hours, plasma drug levels at all doses lower than 600 mg/kg approached or dropped below 1 µg/mL, which is EBO MIC_90_ (Amornchai et al The World Melioidosis Congress 2024). The 600 mg/kg group maintained plasma levels that were equal (1.08 ± 0.97 µg/mL) to the MIC_90_ for EBO. At 24 hours post-dose, plasma concentrations were near the assay’s lower limit of detection, ranging from 0.008 ± 0.002 µg/mL (100 mg/kg) to 0.038 ± 0.018 µg/mL (600 mg/kg; **Table 2**).

**Figure 1:**
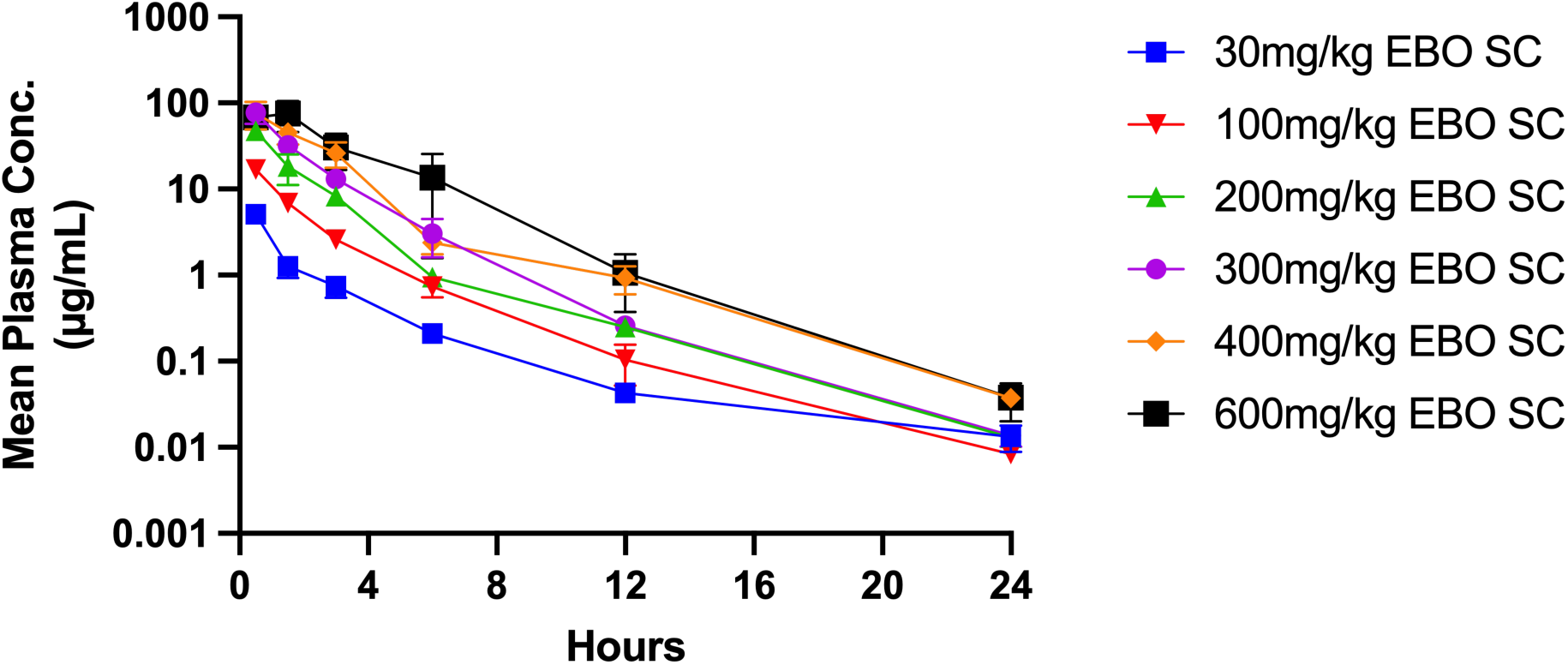
Epetraborole Pharmacokinetic Profile in BALB/c mice. 7-9 week-old BALB/c mice were treated with EBO via subcutaneous injection at doses of 30, 100, 200, 300, 400, and 600 mg/kg. Plasma samples were collected from three mice per time point, and EBO concentrations were quantified using an Agilent 6460 triple quadrupole mass spectrometer. Data represent mean ± SEM.

**Table 2:** Plasma Concentrations and Area Under the Curve (AUC).

| Plasma | 30 mg/kg EBO SC |  | 100 mg/kg EBO SC |  | 200 mg/kg EBO SC |  | 300 mg/kg EBO SC |  | 400 mg/kg EBO SC |  | 600 mg/kg EBO SC |  |
| --- | --- | --- | --- | --- | --- | --- | --- | --- | --- | --- | --- | --- |
| Time (h) | Mean Conc. (µg/mL) | SD | Mean Conc. (µg/mL) | SD | Mean Conc. (µg/mL) | SD | Mean Conc. (µg/mL) | SD | Mean Conc. (µg/mL) | SD | Mean Conc. (µg/mL) | SD |
| 0.5 | 5.098 | 0.467 | 16.841 | 1.508 | 47.606 | 1.148 | 77.532 | 20.593 | 75.982 | 26.49 | 68.228 | 18.95 |
| 1.5 | 1.272 | 0.339 | 6.856 | 0.844 | 18.278 | 7.101 | 32.302 | 4.982 | 45.473 | 12.494 | 75.788 | 29.628 |
| 3 | 0.746 | 0.201 | 2.588 | 0.114 | 8.286 | 0.651 | 13.167 | 2.324 | 26.25 | 8.477 | 30.209 | 13.53 |
| 6 | 0.211 | 0.019 | 0.734 | 0.182 | 9.501 | 1.304 | 3.042 | 1.433 | 2.385 | 0.645 | 13.558 | 11.989 |
| 12 | 0.043 | 0.003 | 0.104 | 0.051 | 0.251 | 0.004 | 0.257 | 0.058 | 0.466 | 0.166 | 1.082 | 0.97 |
| 24 | 0.013 | 0.005 | 0.008 | 0.002 | 0.013 | 0.002 | 0.014 | 0.004 | 0.019 | 0.003 | 0.038 | 0.018 |

| AUC | 30 mg/kg EBO SC | 100 mg/kg EBO SC | 200 mg/kg EBO SC | 300 mg/kg EBO SC | 400 mg/kg EBO SC | 600 mg/kg EBO SC |
| --- | --- | --- | --- | --- | --- | --- |
| Total Area | 7.234 | 27.1 | 110.4 | 124.9 | 168.9 | 267.8 |
| Std. Error | 0.5164 | 1.294 | 7.85 | 12.83 | 22.59 | 54.57 |
| 95% CI | 6.222 to 8.246 | 24.56 to 29.63 | 95.00 to 125.8 | 99.71 to 150.0 | 124.7 to 213.2 | 160.8 to 374.7 |
| AUC:MIC | 7.2 | 27.1 | 110.4 | 124.9 | 168.9 | 267.8 |

To quantify systemic exposure, the area under the curve (AUC₀–₂₄) was calculated for each dose group (**Table 2**). AUC increased with dose, though the relationship was nonlinear. AUC for 30 mg/kg was 7.23 ± 0.52 µg·h/mL, while 600 mg/kg resulted in an AUC of 267.8 ± 54.57 µg·h/mL, with notable inter-animal variability at higher doses. The corresponding 95% confidence intervals (CI) spanned from 6.22–8.25 µg·h/mL at 30 mg/kg to 160.8–374.7 µg·h/mL at 600 mg/kg, highlighting variability in systemic exposure. These findings indicate that EBO is rapidly absorbed, achieves peak plasma levels within 30 minutes, and demonstrates sustained systemic exposure at higher doses. The nonlinear increase in AUC at elevated doses may reflect saturation of clearance pathways. Collectively, these data support dose optimization strategies to maintain effective plasma concentrations and meet pharmacodynamic targets, informing future preclinical and clinical development of EBO for melioidosis.

### Lab reference & clinical strain infection comparison in *Burkholderia* animal model of infection

To evaluate the baseline bacterial burden and disease progression across genetically and geographically diverse *B. pseudomallei* strains, we assessed lung CFU counts 24 hours post-infection in BALB/c mice (**Figure 2**). The use of this mouse strain for *B. pseudomallei* infections to study antibacterial treatments for melioidosis has been previously described [17]. They observed that in BALB/c mice, the pathogen exhibits the same lung, liver, and spleen tropism observed in human acute melioidosis. The comparisons in this study provide context for interpreting strain-specific responses and appropriate strain selection for downstream efficacy studies. Only a few *B. pseudomallei* strains have been used in murine efficacy studies; selecting a diverse panel of strains of clinical origin and known provenance is an important facet of this study for relating the efficacy of EBO in the mouse model to planned treatment of human melioidosis disease. Despite equivalent burdens to establish infection, substantial variability in bacterial loads was observed among the tested strains. Reference strains 1026b and K96243 demonstrated high levels of lung burden, with mean burdens approaching 6.0 log₁₀ CFU. Several clinical isolates, including MSHR435, 406e, NCTC6700, HBPUB10134a, HBPUB10303a, NCTC10274, and isolates 1710a, 1710b, and 1106b, showed moderate growth, with average lung burdens ranging from 5.0 to 5.5 log₁₀ CFU. Notably, strain NCTC10276 exhibited substantially reduced replication, with mean burdens below 3.8 log₁₀ CFU, suggesting diminished *in vivo* fitness or enhanced host containment.

**Figure 2:**
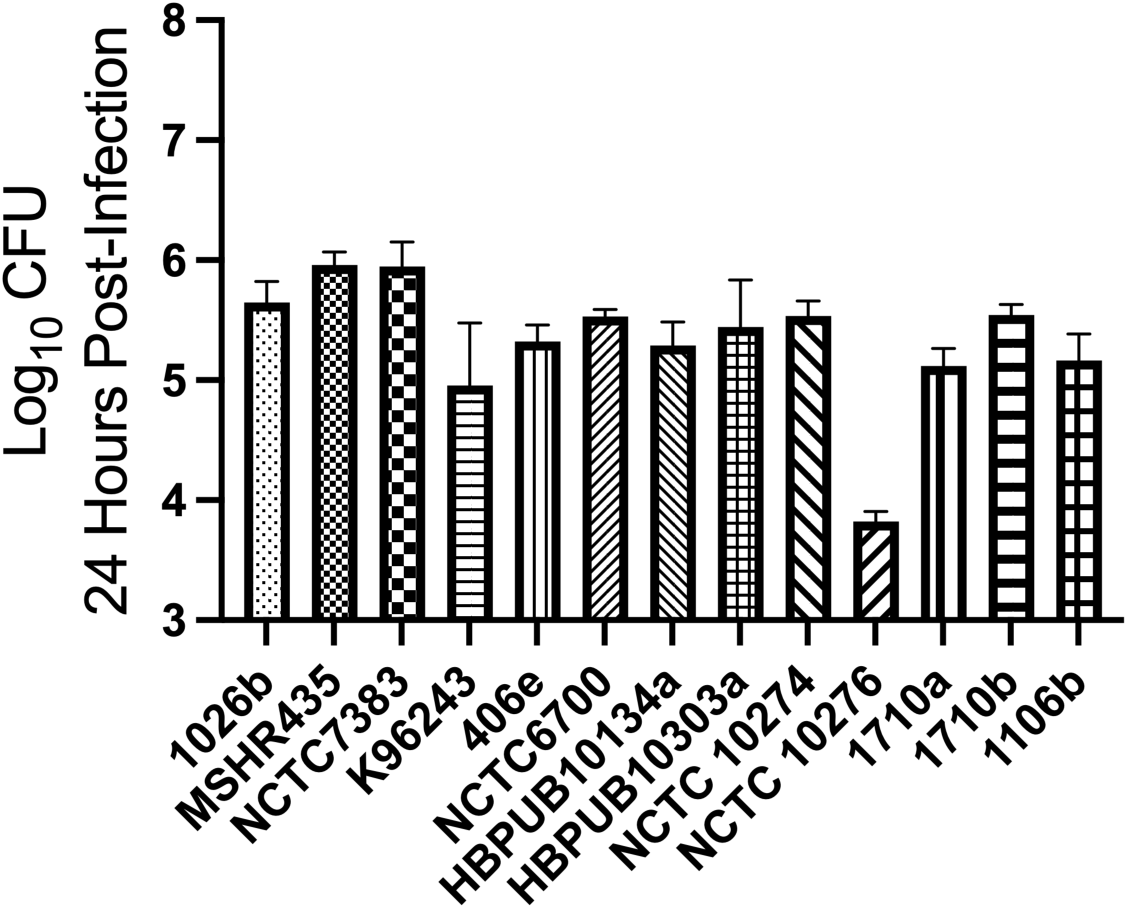
*B. pseudomallei* strain panel characterization in Acute Animal Infection Model. BALB/c mice (7–9 weeks old) were infected intranasally with 5,000 CFU of each *B. pseudomallei* strain. Lung bacterial burdens were quantified at 24 hours post-infection to characterize baseline infection levels prior to treatment initiation in future efficacy studies. Data represent mean ± SEM.

The observed differences in bacterial replication under identical host conditions highlight inherent strain-specific differences, potentially driven by genomic variation, virulence factors, or host-pathogen interactions, and demonstrate the need to use more than 2 laboratory reference strains, 1026b and K96243, for *in vivo* efficacy strains, that have been used for the majority of published murine efficacy studies. In general, the strains progressed during infection, achieving consistent burdens within a narrow range. However, the lesser bacterial burden observed for NCTC10276 emphasizes the importance of including multiple clinically derived strains in efficacy studies to account for biological variability.

These results underscore the need to consider baseline virulence and replication kinetics when evaluating therapeutic outcomes. Strains with higher replication capacity may present greater therapeutic challenges, while those with lower burdens may obscure efficacy signals. Accordingly, strain selection and characterization are critical for generating translationally relevant efficacy data and ensuring that preclinical findings apply to the clinical diversity of melioidosis. To our knowledge, no previous therapeutic study has evaluated efficacy across as many as ten *B. pseudomallei* strains in a single animal model. By systematically profiling *in vivo* bacterial burden across this diverse panel based on clinical isolates, we establish a rigorous foundation for evaluating drug response beyond relying on one or two frequently used reference strains.

### Dose fractionation reveals exposure-dependent efficacy

To evaluate the impact of dose level and dosing frequency on bacterial clearance and to identify the best pharmacodynamic parameter for efficacy, we conducted a dose fractionation study using *B. pseudomallei* strain NCTC7383, the least susceptible isolate to epetraborole in our panel (MIC = 4 µg/mL), which represents the MIC_100_ strain. Mice were treated with varying regimens of EBO or ceftazidime beginning 24 hours post-infection, and lung bacterial burden (log₁₀ CFU) was assessed after 48 hours of therapy (**Figure 3**). The untreated pre-treatment group exhibited a mean lung burden of ∼7.2 log₁₀ CFU. Ceftazidime (300 mg/kg SC TID) significantly reduced this burden to ∼6.2 log₁₀ CFU (p < 0.001), establishing a baseline comparator. In contrast, EBO demonstrated a robust dose-response relationship. Across all EBO treatment groups, bacterial burden reductions ranged from 0.9 to 3.0 log₁₀ CFU (p < 0.001) relative to the pre-treatment control. Notably, the 200 mg/kg SC TID and 100 mg/kg SC TID groups achieved the most substantial reductions, with 3 and 2.9 log₁₀ CFU reductions from the start of therapy, and outperformed ceftazidime by ∼2 log₁₀ CFU. The 33 mg/kg TID dose was the least efficacious, with only a 0.9 log_10_ CFU reduction. The QD groups (600, 300, 100 mg/kg) yielded ∼2.7, 2.0, and 1.6 log₁₀ CFU reductions, respectively, while intermediate BID regimens (300, 150 mg/kg BID) gave ∼2.2 and 1.9 log₁₀ reductions (**Table 3**).

**Figure 3:**
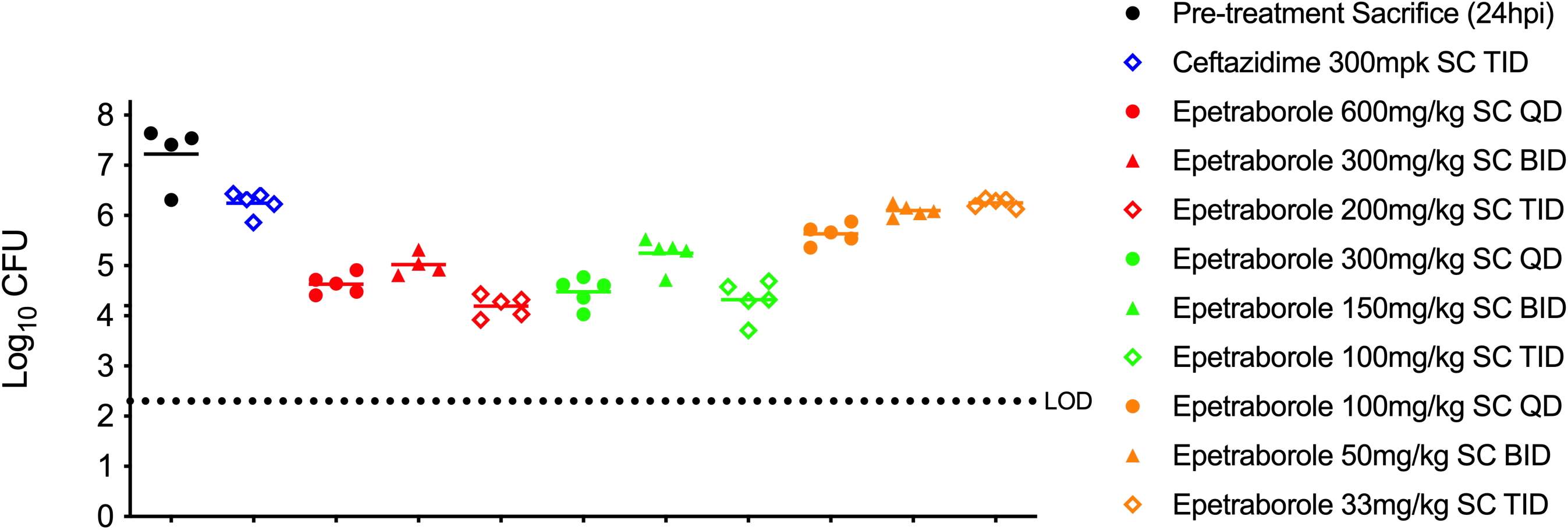
Epetraborole Dose Fractionation. Mice were infected intranasally with 5,000 CFU of *B. pseudomallei* NCTC7383 and treated 24 hours post-infection with EBO subcutaneously in fractionated dosing regimens. Nine treatment groups received total daily doses of 600 mg/kg (QD), 300 mg/kg (BID), 200 mg/kg (TID); 300 mg/kg (QD), 150 mg/kg (BID), 100 mg/kg (TID); and 100 mg/kg (QD), 50 mg/kg (BID), 33 mg/kg (TID). Lung bacterial burdens were assessed 72 hours post-infection (48 hours after treatment initiation). Data represent mean ± SEM, and statistical analysis was done using a two-way ANOVA with Tukey multiple comparisons test (p<0.001=***).

**Table 3:**
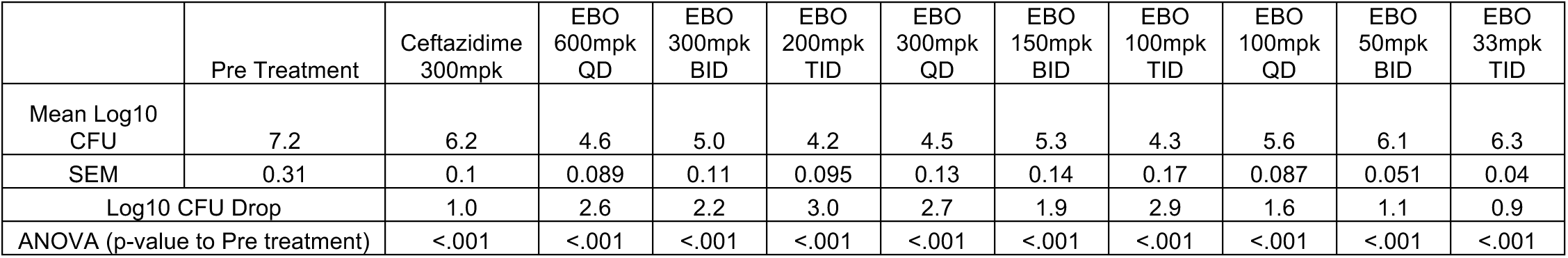
Epetraborole Dose Fractionation Statistics.

To further assess treatment impact, Cohen’s d effect sizes were calculated and categorized as minimal (d < 2.13), moderate (2.13 ≤ d < 3.88), large (3.88 ≤ d < 6.11), or very large (d ≥ 6.11). The 200 mg/kg TID group demonstrated the most pronounced bacterial clearance, with mean and maximum d values of 9.18 and 13.00, respectively. Additional regimens exhibiting large or very large effects included 600 mg/kg QD (mean d = 8.45; max d = 11.5), 300 mg/kg QD (mean d = 6.80), 300 mg/kg BID (mean d = 6.43), 100 mg/kg TID (mean d = 6.18), and 150 mg/kg BID (mean d = 4.27). These results indicate EBO efficacy is driven by total systemic exposure rather than dosing frequency alone. Regimens with higher cumulative daily doses, particularly those administered QD or BID, resulted in greater bacterial reductions, consistent with concentration-dependent pharmacodynamics. Importantly, although NCTC7383 exhibited a higher MIC (4 µg/mL) than laboratory strain 1026b (1 µg/mL), EBO maintained efficacy across dosing strategies, reaffirming its potential even against strains that would cover the entire range of EBO susceptibilities. These data substantiate that EBO is effective against various strains with variable susceptibilities, provides critical insight into dosing strategies, and reinforces the importance of optimizing AUC-driven regimens in future evaluations of epetraborole for melioidosis. Moreover, the study offers an animal PK-PD infection model that can inform preclinical assessment of EBO to support setting *in vitro* susceptibility breakpoints, and dose and dosing selections for future clinical studies of EBO in treating acute melioidosis [18].

### Epetraborole Dose-Response Efficacy Across Different Strains of *B. pseudomallei*

To expand the scope of EBO’s *in vivo* efficacy with other antibacterial agents, we assessed efficacy against a diverse panel of *B. pseudomallei* strains to evaluate strain-specific responsiveness and define optimal dose-response relationships. Ten strains were selected based on prior drug susceptibility and virulence profiling, including reference strains (1026b, K96243) and clinically derived isolates (MSHR435, NCTC7383, 406e, 1710a, 1710b, HBPUB10134a, HBPUB10303a, and 1106b) [9]. Pre-treatment sacrifices confirmed consistent infection establishment, with baseline lung burdens ranging from 5.0 to 6.0 log₁₀ CFU across all strains. Treatment with EBO for 48 hours resulted in dose-dependent bacterial clearance (**Figure 4**). At 200 mg/kg, EBO significantly reduced lung burden in all strains by 1.7–3.7 log₁₀ CFU (p < 0.001), with several animals falling below the limit of detection (LOD = 200 CFU). At 100 mg/kg, 8 of 10 strains showed significant burden reductions (2.2–3.5 log₁₀ CFU), but reductions for MSHR435 and NCTC7383 were not statistically significant. Similarly, 30 mg/kg yielded partial efficacy for 7 of 10 strains (0.9–3.0 log₁₀ CFU reduction), with limited or no effect observed against MSHR435, NCTC7383, and 406e. For reference strain 1026b, EBO showed a clear dose response: 30 mg/kg yielded modest reductions, while 100 mg/kg and 200 mg/kg drove burdens near or below LOD (**Table 4**). Strain-specific susceptibility differences were evident. MSHR435 and NCTC7383 required the highest dose (200 mg/kg) for significant reductions, and even then, reductions were less than those of other strains. These findings align with prior MIC data and underscore the relevance of individual strain characteristics when evaluating therapeutic outcomes.

**Figure 4:**
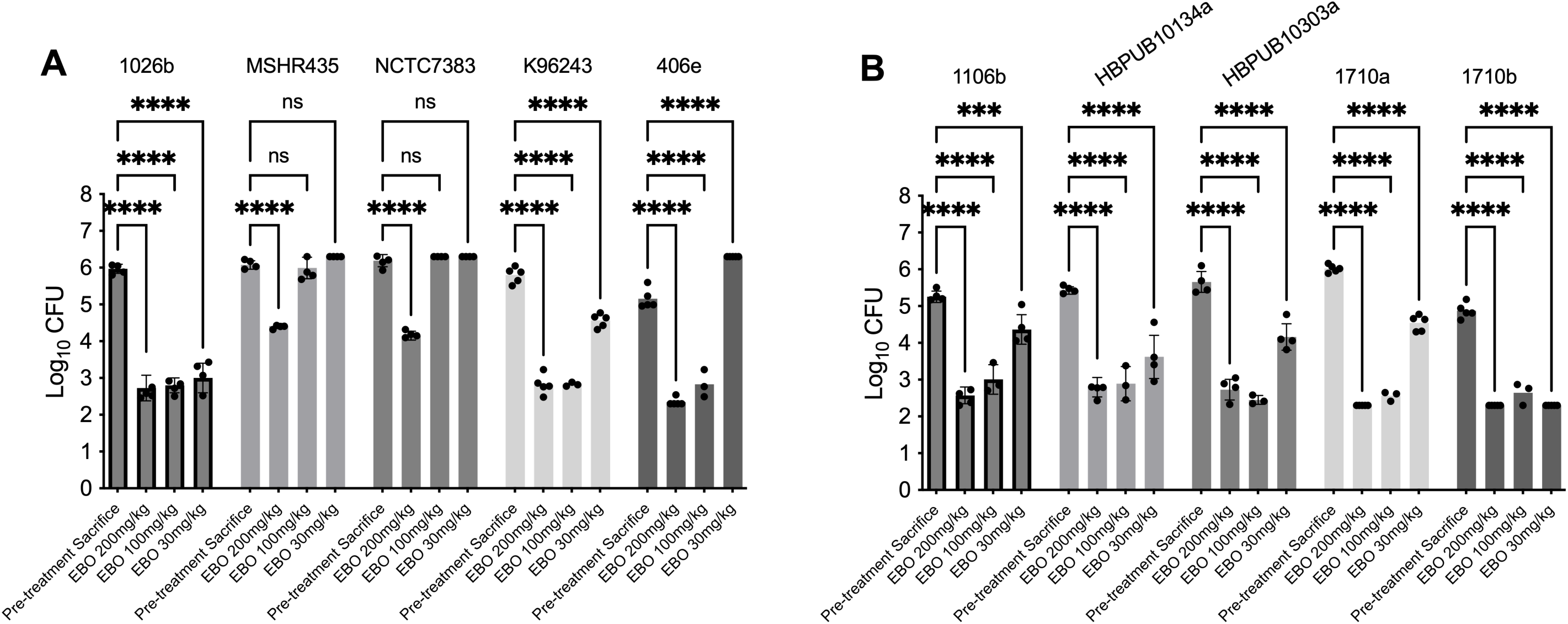
Epetraborole Diverse Strain Panel Efficacy. BALB/c mice (7–9 weeks old) were infected intranasally with 5,000 CFU of diverse *B. pseudomallei* strains. Epetraborole (EBO) was administered subcutaneously once daily at 30, 100, or 200 mg/kg, beginning 24 hours post-infection. Lung bacterial burdens were determined at 72 hours post-infection. Data represent mean ± SEM, and statistical analysis was done using a two-way ANOVA with Tukey multiple comparisons test (p<0.001=***, p<0.0001=****).

**Table 4:**
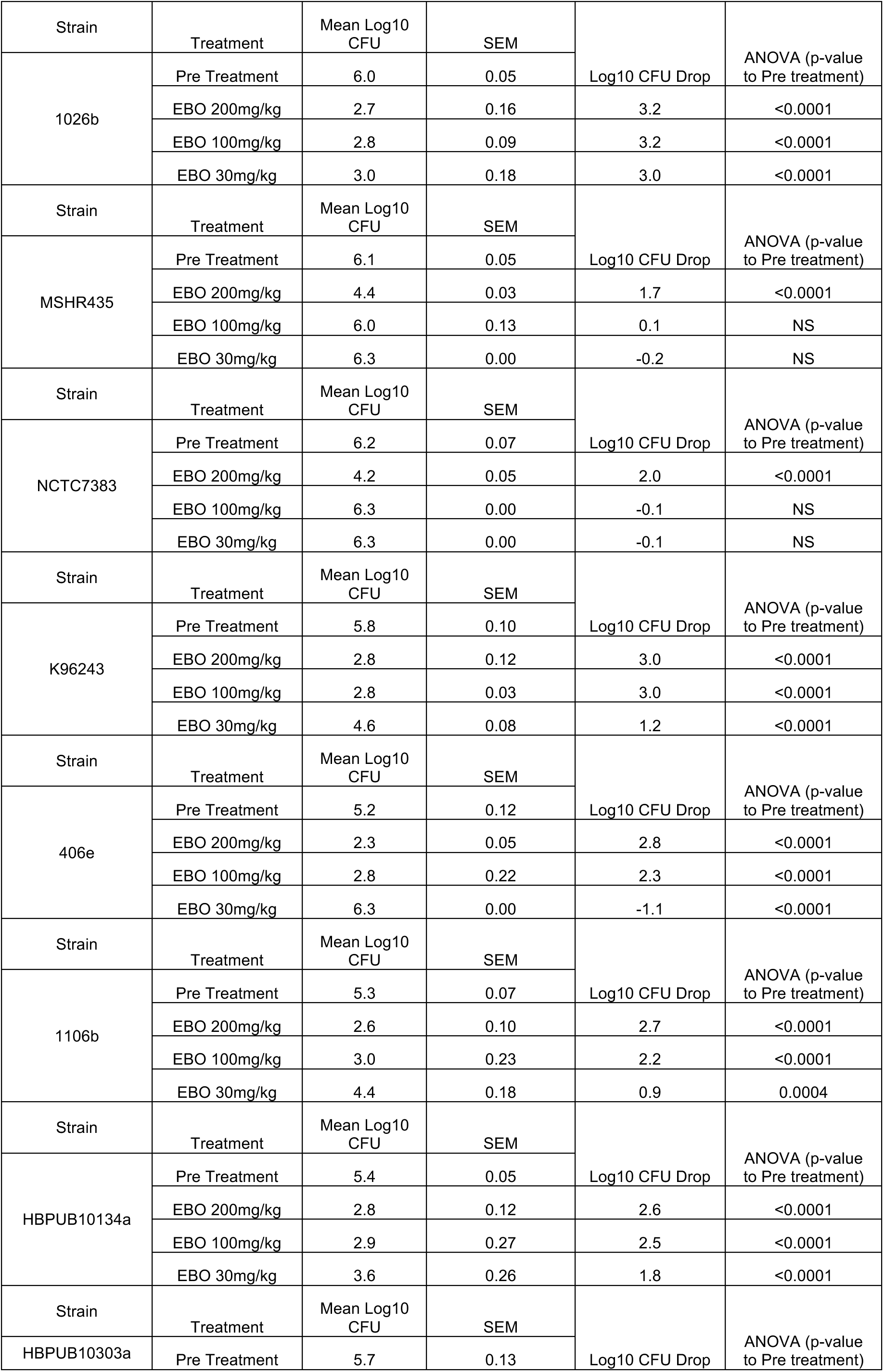
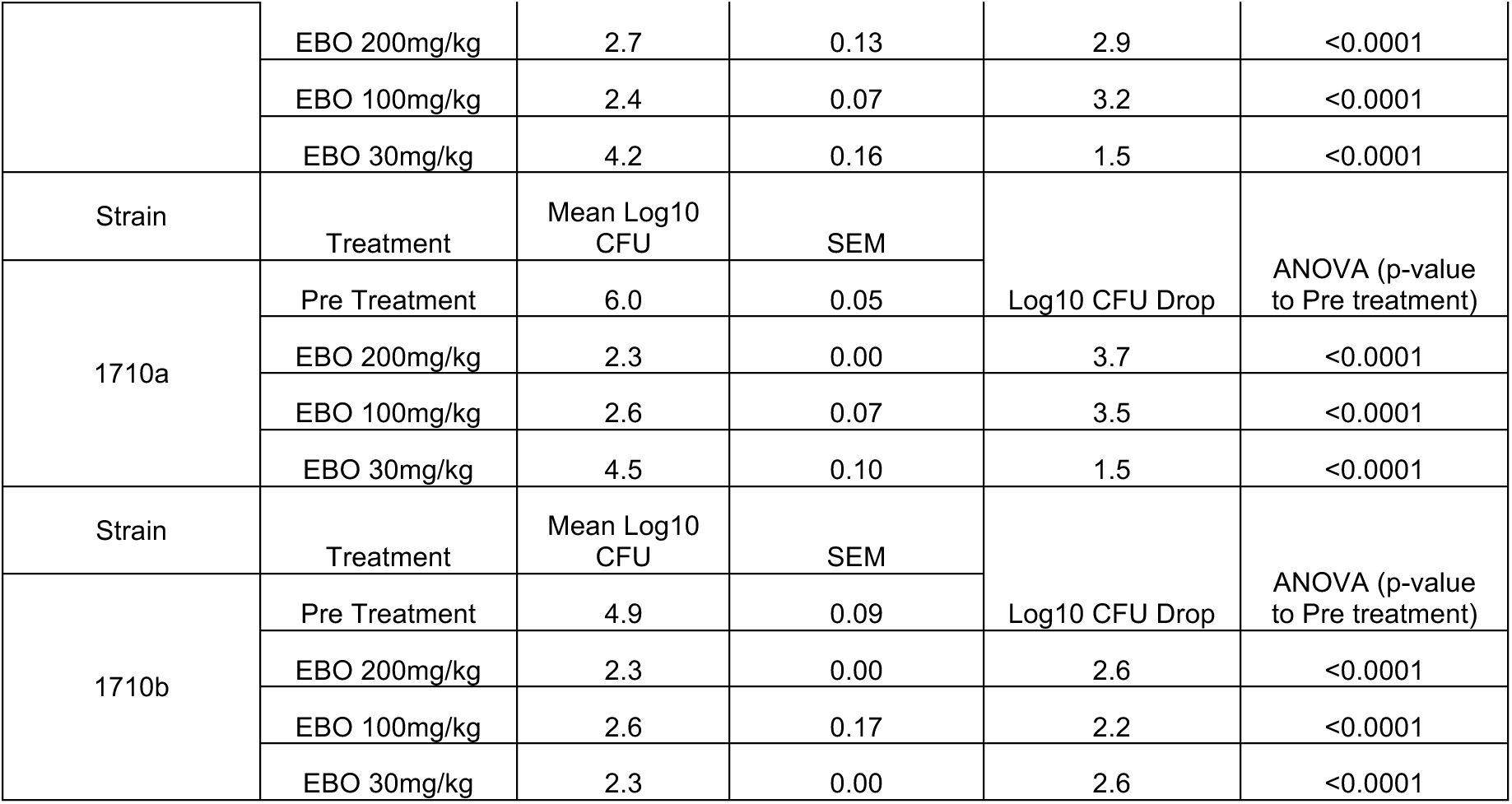
Epetraborole Diverse Strain Panel Statistics.

Effect size analysis using Cohen’s d further quantified the treatment effect. Quartile thresholds were defined as minimal (d < 0.65), moderate (0.65 ≤ d < 5.39), large (5.39 ≤ d < 11.81), and very large (d ≥ 11.81). The 200 mg/kg group demonstrated the strongest mean and maximum effect sizes (mean d = 15.36; max d = 20.20), followed by the 100 mg/kg group (mean d = 10.86; max d = 13.24). While 30 mg/kg showed moderate efficacy (mean d = 9.24; max d = 10.80), it was consistently inferior to higher doses. These findings confirm that EBO achieves dose-dependent reductions in lung bacterial burden across a range of clinically derived *B. pseudomallei* strains, with 8 strains showing greater than 2 log_10_ CFU reductions with 100 mg/kg QD. Only two strains NCTC7383 and MSHR435 show minimal CFU reductions at 100 mg/kg QD, requiring 200 mg/kg QD to achieve greater than 1 log_10_ CFU reduction. This indicates that a minimal dose of 200 mg/kg QD would be necessary to achieve a greater than 1-log_10_ CFU bactericidal kill across all 10 strains tested, and this dose is sufficient to overcome susceptibility differences between strains, emphasizing the importance of dose optimization and susceptibility-guided treatment strategies. These data reinforce the potential of EBO as a broad-spectrum candidate for melioidosis therapy.

## Discussion

Melioidosis presents a significant clinical challenge due to the intrinsic resistance of *B.pseudomallei* and high relapse rates despite prolonged antibiotic therapy [3,4]. Treatment regimens are lengthy, require hospitalization, and often involve multiple antibiotic classes to achieve bacterial clearance [6,7]. In this study, we characterized the pharmacokinetics and *in vivo* efficacy of EBO, a LeuRS inhibitor, and demonstrated its promising activity against clinically derived *B. pseudomallei* strains in a murine acute infection model. Previous work has established a strong link between bacterial burden and treatment outcome, with higher initial loads correlating with reduced therapeutic efficacy and increased relapse risk [14, 19]. Our results showed that EBO maintained relatively stable MICs across a range of inocula similar to ceftazidime and meropenem. These findings suggest that EBO may retain activity in clinical scenarios with high bacterial loads, an important consideration for acute melioidosis, where bacterial burden at presentation can be considerable [3,4]. Pharmacokinetic analysis confirmed rapid absorption and sustained exposure of EBO following subcutaneous dosing, with the AUC:MIC ratio exceeding the target threshold of 30, a threshold associated with targeting bacterial lung infections. The non-linear increase in AUC at higher doses may reflect saturation of metabolic clearance pathways and warrants further study to refine dosing regimens.

EBO exhibited robust *in vivo* efficacy across a diverse panel of clinically derived *B. pseudomallei* strains. Although 100 mg/kg proved effective for most strains, a 200 mg/kg dose was required to reduce bacterial burden by >1 log₁₀ CFU in MSHR435 and NCTC7383—two strains with higher MICs. A 200 mg/kg dose in the satellite PK animal group yields an AUC_0-24_ of 110 µg.h/mL, suggesting that a 2,000 mg IV dose in humans might be required for optimal efficacy against less susceptible strains. However, as previous studies have shown a benefit of adding EBO on top of ceftazidime in mouse infection models of melioidosis [15], a lower exposure of EBO might be equally efficacious in combination. These findings are consistent with strain-specific variability in antimicrobial susceptibility, likely influenced by metabolic adaptations, efflux mechanisms, or biofilm formation [9,11,12].

The dose fractionation study provided additional insight into EBO’s pharmacodynamics. Regimens delivering the same total daily dose but differing in frequency (QD, BID, TID) achieved comparable reductions in bacterial burden, indicating that efficacy is exposure-driven rather than time-dependent. This aligns with established models for other concentration-dependent agents such as aminoglycosides and fluoroquinolones [17]. Importantly, less frequent dosing could simplify administration schedules and enhance treatment adherence, particularly in resource-limited or outpatient settings [7].

Pharmacodynamic relevance was assessed using the free AUC:MIC ratio, a key parameter for concentration-dependent antimicrobials. Although for Gram-negative pathogens, an AUC:MIC >125 is associated with optimal efficacy [16,17], EBO has been shown to require a free-drug plasma AUC:MIC of 23.8 for net bacterial stasis in a murine *P. aeruginosa* lung infection model, where 92% of EBO is unbound in mouse plasma [20]. EBO achieved this target at doses ≥200 mg/kg, but yielded at least a 1.7-log_10_ CFU kill rather than bacterial stasis, which might suggest that *B. pseudomallei* is more susceptible to EBO than *P. aeruginosa*. This could be because EBO has a 10-fold higher concentration in alveolar macrophages than epithelial lining fluid in mice, as in humans (Tenero et al. 2013 57:3334), and *B. pseudomallei* is a facultative intracellular pathogen.

The ability of EBO to achieve significant bacterial reductions across lab and clinical strains supports its potential for use as a frontline or adjunctive therapy. The PK-PD results from this study and the observation that there is good concordance between such animal studies and data from human infections [18] indicate that an effective human equivalent dose of 2000 mg IV (q24h), which yields an AUC_0-24_ of 107 µg.h/mL, would offer adjunctive treatment (i.e., with current standard of care antibacterials) of benefit for patients with acute melioidosis [18], noting that adding EBO on top of ceftazidime was at least additive in mouse models, and a lower might also be possible (reference previous EBO Bpm paper). Given the high relapse rates reported for melioidosis even after prolonged treatment [3], EBO’s rapid bactericidal activity and distinct mechanism of action may help shorten therapy duration and improve long-term outcomes. Our findings also suggest its suitability for combination therapy, especially when paired with standard-of-care agents to enhance bacterial killing and reduce the risk of resistance [13].

EBO targets LeuRS, a novel mechanism of action among agents tested for melioidosis. Unlike ceftazidime and meropenem, which inhibit cell wall biosynthesis, epetraborole disrupts protein synthesis, offering a complementary mechanism with potential synergy and reduced cross-resistance risk [13]. This is particularly relevant as reports of multidrug-resistant *B. pseudomallei* are increasing, especially in regions with high antibiotic pressure [9]. EBO’s distinct mechanism and efficacy profile make it a promising candidate for further preclinical development and eventual clinical evaluation.

## Conclusion

This study demonstrates the potential of EBO as a therapeutic candidate for treating melioidosis. EBO exhibited potent, dose-dependent *in vivo* efficacy across multiple *B. pseudomallei* strains, including clinical isolates, and substantially reduced bacterial burden. Its favorable pharmacokinetic profile, including rapid absorption and sustained systemic exposure, supports a concentration-dependent mechanism of action consistent with effective antimicrobial therapy. Importantly, EBO maintained activity even in strains with reduced susceptibility and demonstrated efficacy at high bacterial loads at 200 mg/kg doses, which is possible to achieve with a human dose of 2,000 mg IV q24h. These features position EBO as a viable candidate for melioidosis therapy. This is the first *in vivo* melioidosis treatment study to demonstrate efficacy across such a broad strain panel, addressing an important gap in preclinical testing. Including ten genetically diverse clinical isolates affirms EBO’s potential as a broad-spectrum agent capable of overcoming the challenge of strain-specific variability in therapeutic response. Together, these findings support the continued development of EBO as a novel antimicrobial agent that addresses key limitations of current melioidosis treatments and offers a promising path forward in improving patient outcomes for this neglected and often lethal infectious disease.

## Acknowledgment

M. Nurul Islam provided experimental assistance with the drug analyses. We want to thank Lab Animal Resources (Colorado State University) for the outstanding care of the animals used in these studies. The authors have no conflict of interest to declare. The data supporting this study’s findings are available upon reasonable request.

## Transparency declarations

All other authors: none to declare.

## Author contributions

M.R.K.A. and R.A.S. defined the overall study objectives. J. E. C. and V. G. performed the determination of modal MIC and animal studies. D. F. and G. D. performed mass spectrometry and PK. J. E. C. and R. A. S. performed writing.

